# A high-quality reference genome and tissue expression atlas for the European lobster (*Homarus gammarus*)

**DOI:** 10.1101/2025.09.29.677193

**Authors:** Josephine R Paris, Tom L Jenkins, Joan Ferrer Obiol, Manu K Gundappa, Tim Regan, Lahcen Campbell, Gareth L Maslen, Jorge Alvarez-Jarreta, Sarah Dyer, Aaron R Jeffries, Georgina Murray, Audrey Farbos, Lisa K Bickley, Bas Verbruggen, Kelly S Bateman, Carly L Daniels, Charlie D Ellis, Thomas J Ashton, Charles R Tyler, Grant D Stentiford, Ross Houston, Tim P Bean, Ronny van Aerle, Daniel J Macqueen, Jamie R Stevens, Eduarda M Santos

## Abstract

The European lobster (*Homarus gammarus*) and its sister species, the American lobster (*Homarus americanus*), are notable for their remarkable immunity and longevity, with lifespans reaching up to 80 years in the wild. A reference genome is available for the American lobster, but not yet for the European lobster, despite its ecological significance, and importance to fisheries and aquaculture. Here, we present a high-quality genome assembly and annotation for the European lobster. The assembly spans 1.76 Gb, with a scaffold N50 of 1.82 Mb and a BUSCO completeness of 97.6%. As observed in the American lobster, the total assembly span is substantially smaller than genome size estimates derived from flow cytometry, performed using independently sampled European lobsters (3.18 - 3.42 Gb). This discrepancy may reflect the highly repetitive nature of decapod genomes, with 51.8% of the *H. gammarus* assembly consisting of repetitive elements. Leveraging a comprehensive multi-tissue RNA-seq dataset, we annotated 23,223 protein-coding genes and characterised gene expression across ten tissues to generate a tissue-level gene expression atlas, available at www.LobsterGeneX.com. Using single-copy orthologs, we estimated a divergence time of 26 Mya (95% HPD 22 - 30 Mya) between *H. gammarus* and *H. americanus*, corresponding to the Oligocene-Miocene boundary. We also identified *Homarus*-specific gene duplications with roles in immunity and longevity, including telomere maintenance. The reported genomic resources can facilitate future research into lobster biology, support sustainable fisheries and aquaculture management practices, and enable investigations of the evolutionary mechanisms underlying basic biological processes, notably immunity and longevity in Homarid lobsters.

**Significance Statement:** The European lobster (*Homarus gammarus*) and the American lobster (*Homarus americanus*) are large benthic decapod crustaceans with significant seafood value, known for their remarkable longevity, with typical lifespans of 30-55 years in the wild and maximum lifespans of up to 80 years. Lobsters grow, reproduce and regenerate limbs throughout their life, and there are very few reports of tumours or age-related diseases. We generated and herein share a high-quality genome assembly and annotation for the European lobster alongside a tissue expression atlas, <u>LobsterGeneX</u>, which enables the visualisation of gene expression profiles across ten tissue types. Based on the genetic information generated, we also provide an estimated time for the divergence between *H. gammarus* and *H. americanus* (26 Mya) and identify duplication events for genes related to immunity and longevity. The European lobster reference genome will facilitate further research for investigations into local adaptation in lobster populations, genomic mixing in Homarid hybrids, and identification of genes of interest in aquaculture, ageing, regeneration, disease and cancer resistance.

## Introduction

Decapods are a diverse and widespread group of crustaceans, with many species playing important roles in ecosystem functioning and/or possessing high commercial value (Behringer & Duermit-Moreau 2021). The characteristics and content of decapod genomes are notoriously variable. For instance, genome assembly sizes range from 0.15 Gb (*Pandalus platyceros*) to 5.86 Gb (*Palaemon carinicauda*), and haploid chromosome numbers range from 8 (*Trichoniscus pusillus*) to 104 (*Paralithodes camtschaticus*) (Challis et al. 2023). A particularly notable feature of sequenced decapod genomes is that the assembled genome is typically shorter than estimates derived from *k*-mer profiling and/or flow cytometry. This discrepancy has been largely associated with the highly repetitive nature of decapod genomes (Rutz et al. 2023; Yuan et al. 2023; Zeng et al. 2025) but may also be due to somatic endopolyploidy (Lécher et al. 1995; Korpelainen et al. 1997), and supernumerary B chromosomes (Lécher et al. 1995). These characteristics highlight the complexity of decapod genomes and the importance of developing high-quality reference assemblies to better understand their genome architecture and evolution, as well as for the provision of genomic/genetic resources for biological and functional studies more generally.

The European lobster (*Homarus gammarus*) is a large decapod crustacean of considerable economic and ecological importance. It is a highly valued seafood species with a long history of fisheries exploitation across the northeast Atlantic and Mediterranean coasts. However, slow growth rates and size-specific fecundity (Ellis, Knott, et al. 2015) have increased its vulnerability to overfishing and stock collapse (Kleiven et al. 2022). This has led to fishing restrictions and numerous efforts across Europe to supplement wild stocks with hatchery-reared juveniles to mitigate declines and maintain population sizes (Jenkins et al. 2020; Ellis, Hodgson, et al. 2015). Recent progress in rearing techniques has shown promise for developing lobster aquaculture (Clarke et al. 2023; Halswell et al. 2018; Hinchcliffe et al. 2022), which may help alleviate current fisheries pressures and unlock new markets for lobster trade. European lobsters also have many interesting biological features. For example, when undisturbed by fisheries, they have an average lifespan of 30-55 years and can reach a maximum lifespan of 80 years (Sheehy et al. 1999). They continue to grow, reproduce and regenerate limbs throughout life with no observable decline in function (Vogt 2012), and very few reports of tumours or ageing diseases have been reported, despite their marked longevity (Vogt 2008).

While a reference genome is available for the American lobster (*Homarus americanus*) (Polinski et al. 2021), the only other extant *Homarus* species, no such resource currently exists for the European lobster. Given its biological, fisheries, and aquaculture importance, a reference genome would represent a valuable resource for the research community, for example, by facilitating investigations into local adaptation (Jenkins, Ellis, Triantafyllidis, et al. 2019; Ellis et al. 2023, 2024), for characterising the genetic basis of interspecific hybridisation (Ellis et al. 2020), and aiding the identification of genes associated with desirable traits in aquaculture (Houston et al. 2020; Hinchcliffe et al. 2022). Here, we address this deficit by reporting a high-quality reference genome and annotation for the European lobster. We use the presented genome and its annotation to: (i) quantify gene expression across ten tissue types; (ii) estimate the divergence time between *H. gammarus* and *H. americanus*; and (ii) survey *Homarus* genes linked to immunity and longevity.

## Results and Discussion

### Genome assembly, repeat content, and annotation

The European lobster assembly presented here consists of 4,133 scaffolds (4,350 contigs) and spans 1.76 Gb, with a scaffold N50 of 1.82 Mb (contig N50 = 1.6 Mb). BUSCO analysis indicates a high completeness of 97.6% (arthropoda_odb10, *n* = 1013), which is similar to the BUSCO completeness of *H. americanus* (96.4%). Karyotyping of both *H. gammarus* and *H. americanus*, and crustacean genomes in general, has identified a highly variable number of chromosomes, suggesting that supernumerary chromosomes exist in both species (Hughes 1982). To address this, we performed an analysis of ploidy using Smudgeplot (Supplementary Figure S1). This indicated a diploid genome with the majority of *k*-mer pairs (92%) assigned to an AB configuration with a minor AAAB signal (8%), which may reflect local copy number variation, aneuploidy, or the presence of supernumerary chromosomes. In the European lobster, meiotic counts vary from *n* = 45 - 88. Taking meiotic counts as an approximation for the number of chromosomes, ∼16% of the assembly is contained in the largest 45 contigs and ∼25% of the assembly is contained within the largest 88 contigs. This demonstrates that although the genome assembly is high-quality, future attempts to scaffold these contigs into chromosomes using chromosome-conformation capture techniques would be useful for increasing contiguity. Nevertheless, the overall contig count (4,350) is more than 10-fold lower than the American lobster assembly (55,595). For the European lobster, we estimated a *k*-mer quality value (QV) of 38.06 and a *k*-mer completeness of 89.16%. Similarly, the American lobster had a *k*-mer QV of 22.59 but a *k*-mer completeness of only 17.32%. A full comparison between the assembly statistics of the European lobster and American lobster can be found in Table 1.

**Table 1.**
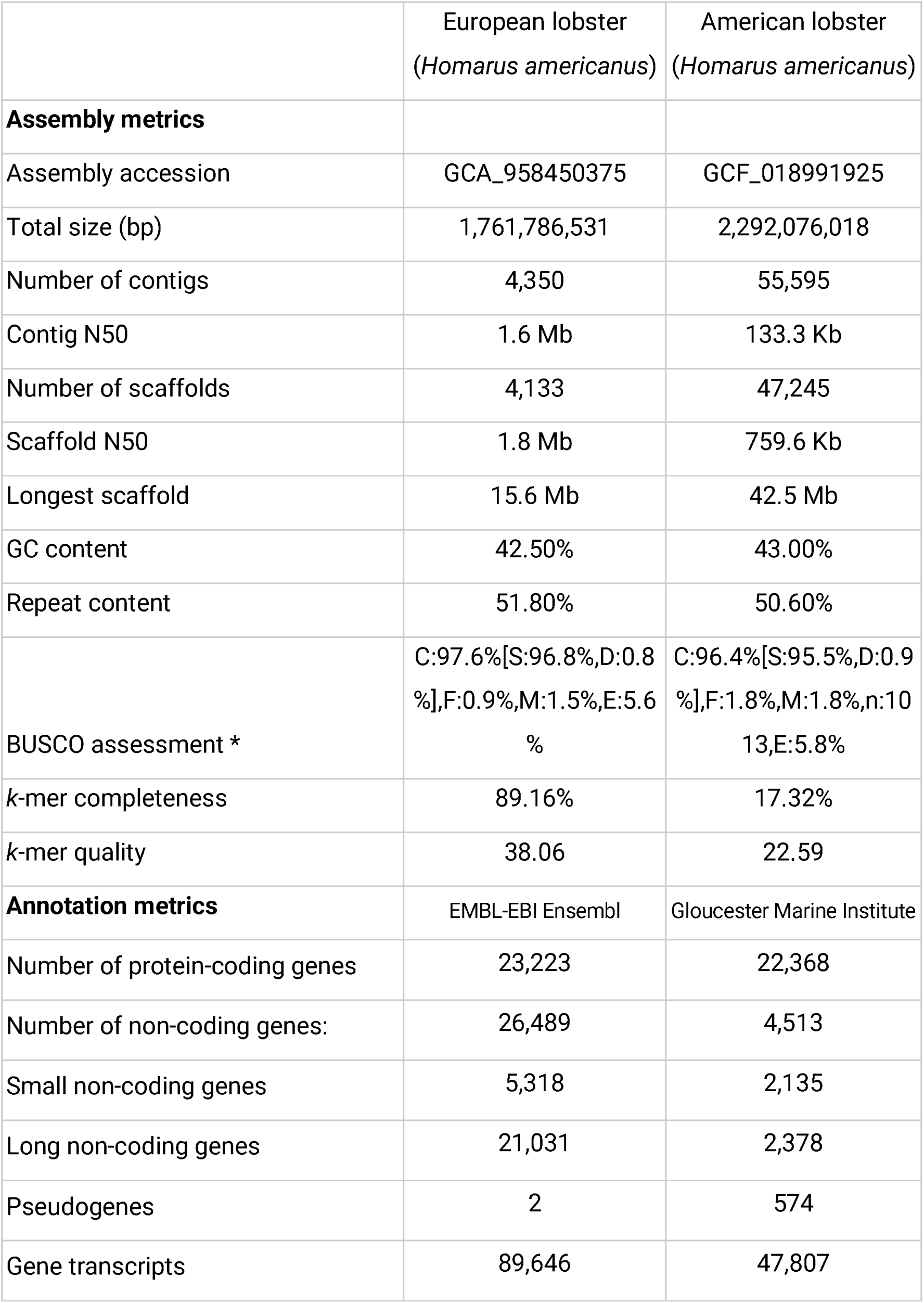
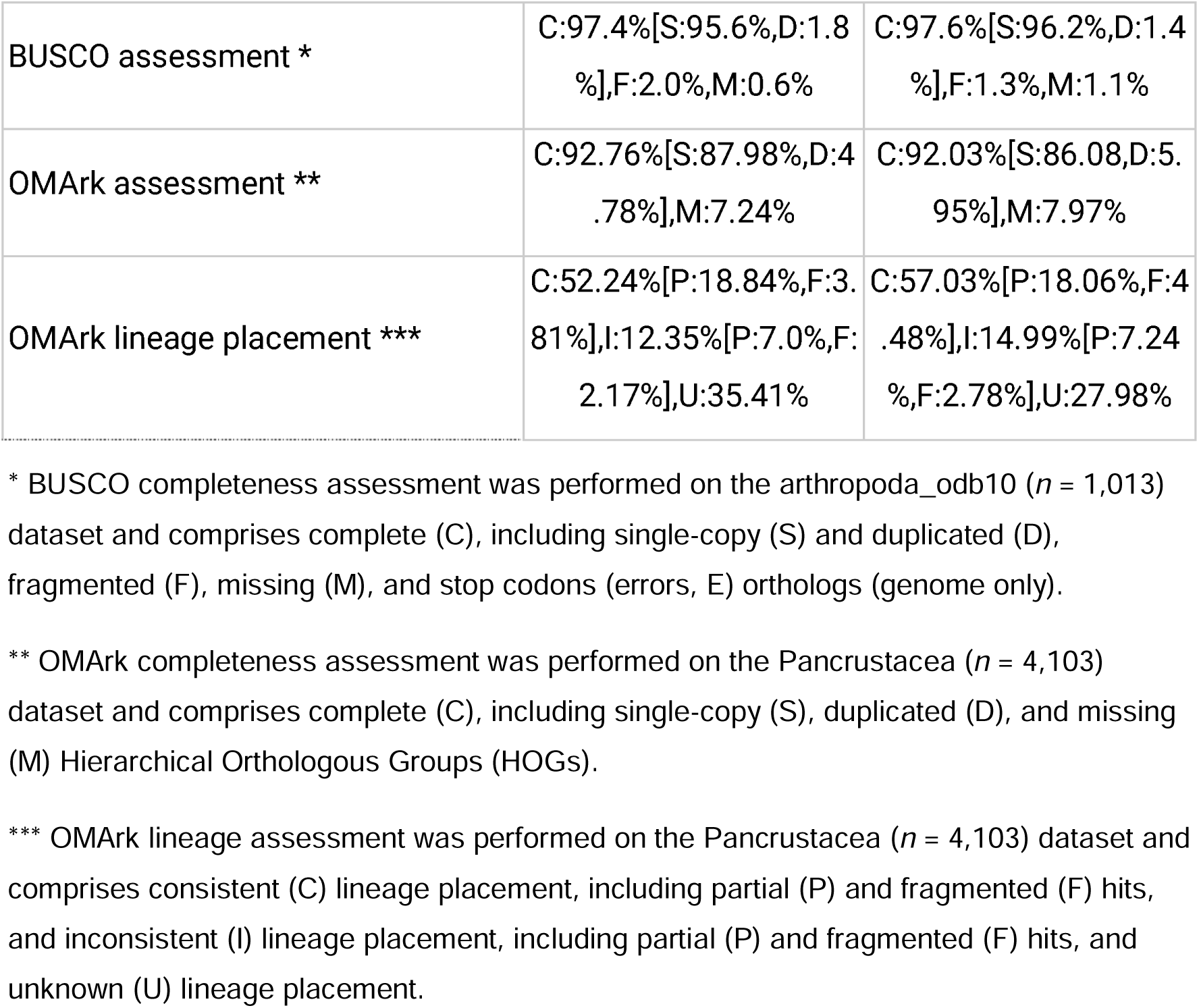
Comparison of genome assembly and annotation statistics for the European lobster (*Homarus gammarus*) and American lobster (*Homarus americanus*).

As an example of how the quality of the assembly might facilitate future functional studies, we examined the Down syndrome cell adhesion molecule (*Dscam*) gene, a member of the immunoglobulin superfamily with roles in neural development and immune function (Schmucker et al. 2000; Ng & Kurtz 2020). In arthropods, *Dscam* produces multiple pathogen specific receptors via immune responsive alternative splicing, generating molecular complexity analogous to vertebrate antibodies (Li et al. 2022). In the American lobster genome, *Dscam* was fragmented across two scaffolds, preventing complete characterisation (Polinski et al. 2021). By contrast, in *H. gammarus* we identified a single, intact *Dscam* locus (ENSHGAG00000010168) on scaffold 3867 (Supplementary Figure S2), comprising 1,139 amino acids encoded by 23 exons (22 coding). *Dscam* expression was detected across all tissues, with the highest levels in neural tissue followed by the heart (Supplementary Figure S3). The recovery of a full-length *Dscam* locus highlights the utility of the European lobster 1 genome for investigating gene function and evolution, particularly for complex and immunologically relevant genes.

Genome size estimation using *k-*mer profiling (21-mers) initially produced an estimated genome size of 2.04 Gb. However, the estimate decreased to 1.80 Gb when reducing the number of repeat *k-*mers included in the estimate, which is comparable to the assembly span. Indeed, analysis of repeat content revealed that 51.8% of the assembly comprises repetitive elements (Figure 1b). Similarly, re-analysis of the American lobster repeat landscape revealed that 50.6% of the assembly comprises repetitive elements. Total repeat content and transposable element (TE) families were very similar between the two species (Supplementary Table S1). Likewise, a recent comparative analysis of TEs across multiple crustaceans identified moderate levels of TE diversification between *H. americanus* and several other species (Zeng et al. 2025). Finally, flow cytometry, performed with three independently sampled individuals, provided an estimated a genome size of 3.18 - 3.42 Gb for *H. gammarus*, while similar estimates for *H. americanus* are 4.30 - 4.65 Gb, nearly double the size of the assembly span in both cases. Given that both assemblies show a high BUSCO completeness but a lower *k*-mer completeness, this discrepancy is likely explained by the highly repetitive nature of these genomes and suggests that both *Homarus* lobster assemblies are missing or have over-collapsed highly repetitive regions.

**Figure 1.**
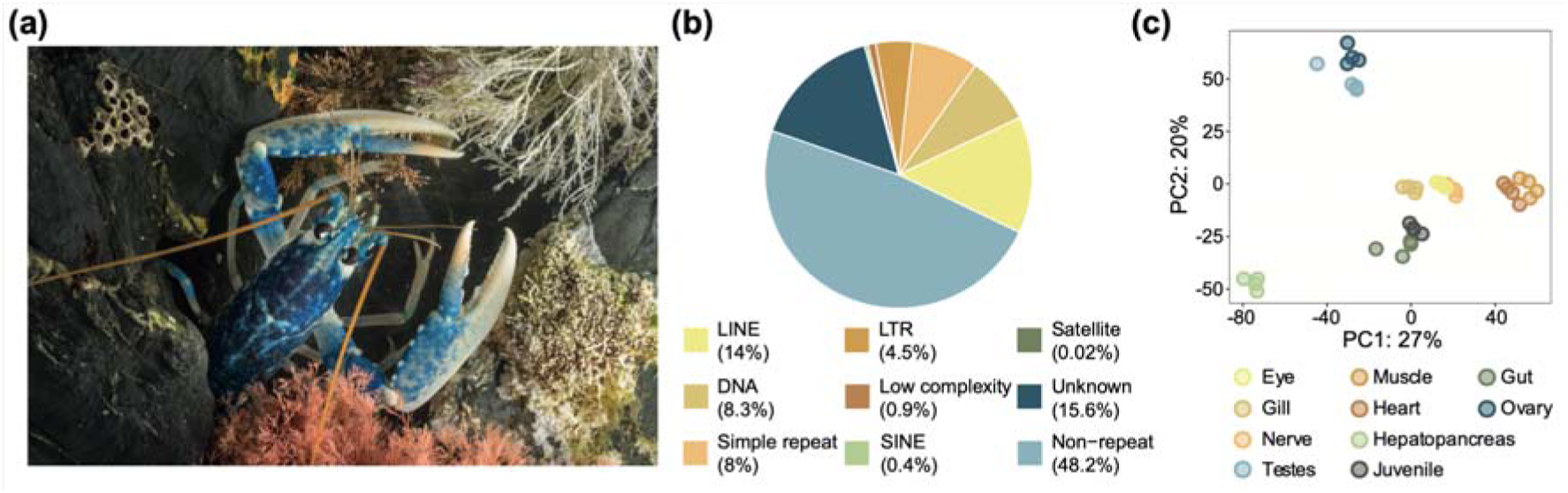
Genome assembly and annotation of the European lobster, *Homarus gammarus* (**a**) Photograph of a juvenile European lobster (photograph credit: Alex Hyde). (**b**) Characterisation of the transposable element (TE) landscape, showing that >50% of the genome is repetitive. (**c**) Principal Component Analysis (PCA) of normalised gene expression counts across the 10 tissues used in the LobsterGeneX (www.LobsterGeneX.com) gene expression atlas.

For structural and functional genome annotation, we generated a comprehensive RNA-seq dataset from ten tissues (eye, gill, gut, heart, hepatopancreas, juvenile, muscle, nerve, ovary, testes). The EMBL-EBI Ensembl Gene Annotation pipeline predicted 23,223 protein-coding genes, (available on Ensembl Metazoa; https://metazoa.ensembl.org/Homarus_gammarus_gca958450375v1). BUSCO and OMArk analysis revealed high annotation completeness (BUSCO completeness: 97.5%; OMArk completeness: 92.76%). Within OMArk, 52.24% of the proteome showed consistent lineage placement, 12.35% showed inconsistent lineage placement, and 35.41% showed an unknown placement. No proteome contamination was identified. Similar OMArk results were obtained using the RefSeq annotation of the American lobster genome annotation (Table 1) suggesting a high number of novel protein-coding genes. Compared to the American lobster annotation performed using a different annotation pipeline, we identified a higher number of long non-coding RNA (IncRNA) genes. Whilst some of these lncRNAs may reflect genuine biological transcripts and represent an interesting avenue for future research (Kim et al. 2017), the apparent inflation in lncRNAs largely reflects technical artefacts. These typically arise from variability in transcript alignments and from the failure to properly collapse alternative splice isoforms, problems that are further exacerbated when gene models are trained on scaffold-level genome assemblies. This highlights the importance of standardising annotation pipelines (Prieto-Baños et al. 2025).

### Tissue-expression atlas for the European lobster

Our multi-tissue RNA-seq data showed a distinct pattern of expression differences between tissues (Figure 1c). Inspired by the growing number of tissue and cell gene expression atlases (Freeman et al. 2012; Papatheodorou et al. 2018; Papatheodorou et al. 2020), we used our RNA-seq dataset to develop a tissue expression atlas for all 23,223 protein-coding genes in the European lobster genome. The atlas (LobsterGeneX; available at www.LobsterGeneX.com) is an interactive web application which allows users to select a gene and visualise its normalised expression across the 10 tissues with four biological replicates. We envisage this open resource will be informative for future studies aiming to characterise the activity of lobster genes, for example those related to immunity and longevity, and the optimisation of hatchery practices.

### Divergence time between *H. gammarus* and *H. americanus*

A phylogeny built using 1295 orthologs from nine decapod species showed the expected evolutionary relationships among the included species. Using the 25 most clock-like single-copy orthologs, and three calibration points, we time-calibrated the phylogeny and inferred a divergence time of 26 Mya (95% HPD 22–30 Mya) between *H. gammarus* and *H. americanus* (Figure 2). Results were robust to different clock models and changes in the prior distributions of calibration points (Supplementary Table S2). This divergence time coincides with the Oligocene-Miocene boundary, and the establishment of the Antarctic Circumpolar Current (ACC), which resulted in a large surface water productivity turnover, accompanied by the upwelling of nutrient-rich waters and changes in global ocean circulation (Rodrigues de Faria et al. 2024; Villa et al. 2014). Our estimated divergence time is markedly more recent than an estimate (100.9 Mya) derived from three mitochondrial genes (16S, 12S, COI) and three nuclear genes (18S, 28S, H3) (Bracken-Grissom & Ahyong 2014) and is slightly older than an estimate derived from 16S mtDNA, 18S and 28S rRNA, and the histone H3 gene (18 Mya; Porter et al. 2005).

**Figure 2.**
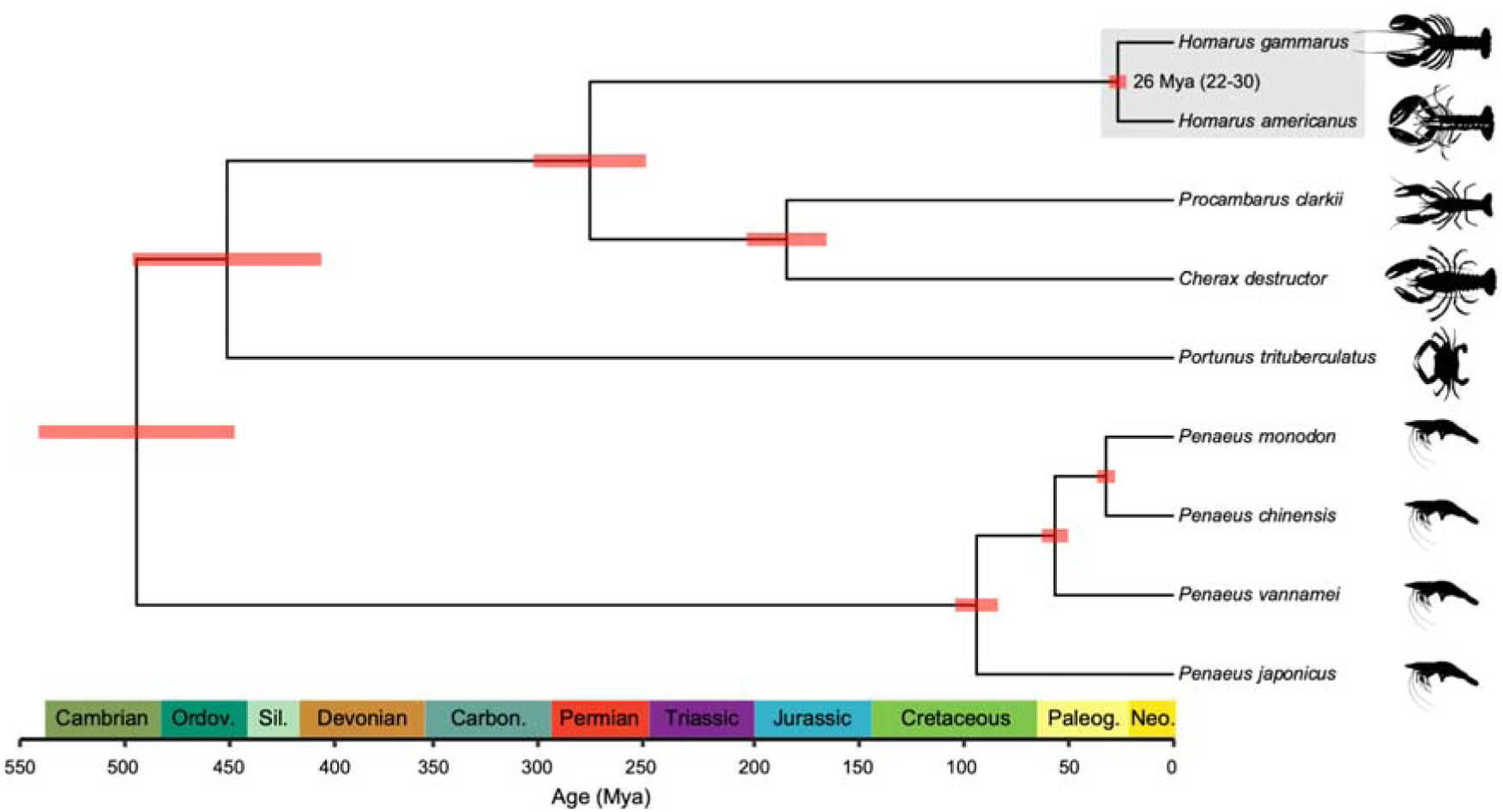
Divergence dating between the European lobster (*Homarus gammarus*) and the American lobster (*Homarus americanus*). BEAST time-tree of nine selected decapod species generated using 25 clock-like orthologs. The grey box highlights the estimated divergence time between *H. gammarus* and *H. americanus* (26 Mya; 95% HPD: 22-30 Mya) and the red horizontal node bars show the 95% HPD for node ages.

### Gene duplications in *Homarus*

In a comparison to seven other decapod species, we identified eight simple duplications (Figure 3a; Supplementary Table S3) and 80 complex duplications (Figure 3b; Supplementary Table S4) in the *Homarus* branch. We define simple duplications as cases where a gene is present as a single copy in all other decapods but appears in two copies in both *Homarus* species, whereas complex duplications occur in gene families that are multicopy across decapods but show further expansion in *Homarus* (see Methods for full statistical treatment).

**Figure 3.**
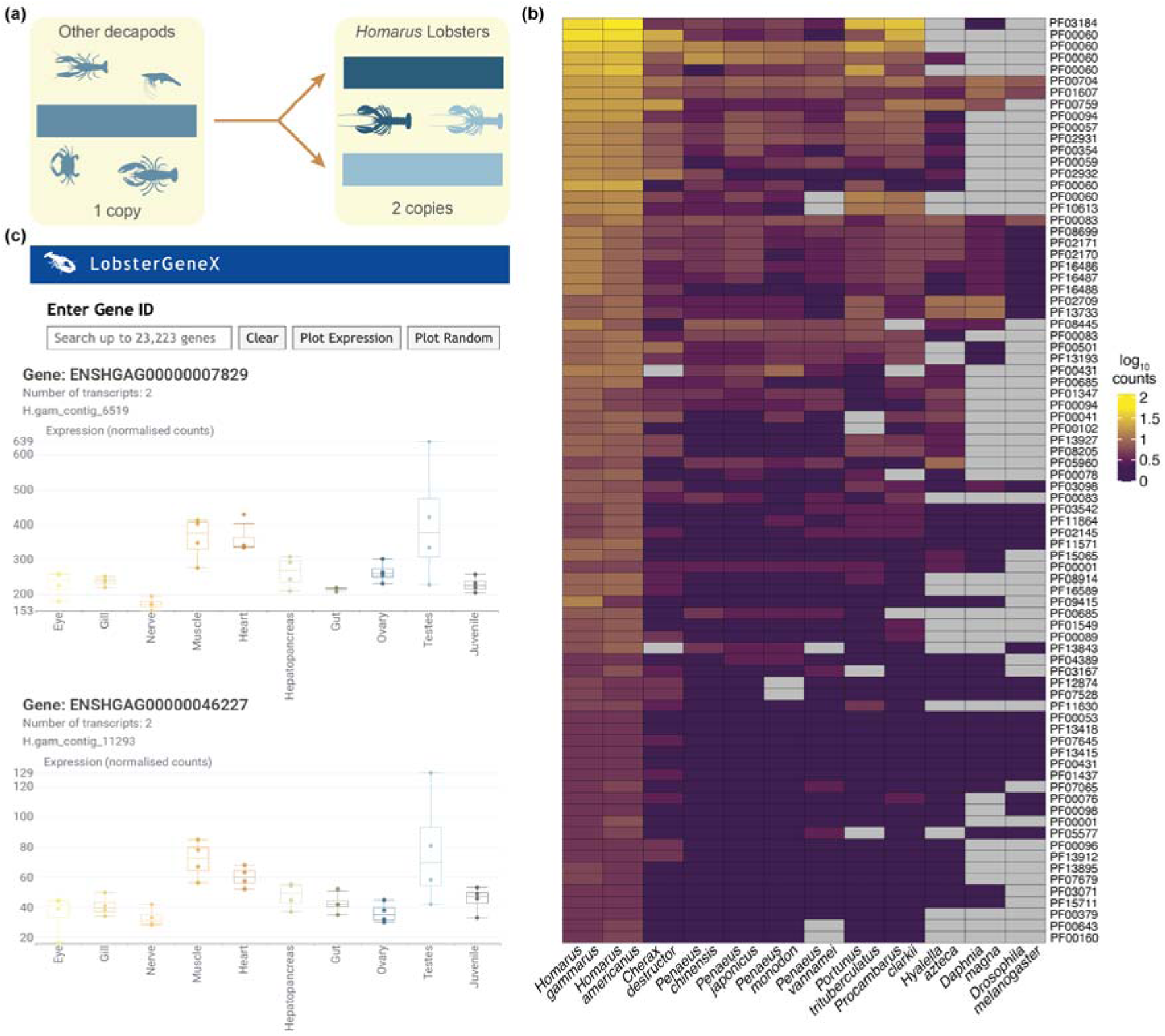
Gene duplications in *Homarus*. (**a**) Schematic representation of identification of the simple duplications unique to the *Homarus* branch. (**b**) Heatmap showing the log_10_ normalised counts of the complex duplications across the two *Homarus* species, the European lobster (*Homarus gammarus*) and the American lobster (*Homarus americanus*) compared to the nine other decapod species and three outgroups. (**c**) Visualisation of the gene expression of the *est1* simple duplication (gene IDs: ENSHGAG0000007829 and ENSHGAG0000046227) in LobsterGeneX. These genes have the Pfam domains PF10373 & PF10374, which have the Pfam descriptions: *est1* DNA/RNA binding domain and Telomerase activating protein *est1*, respectively.

Among the expanded families, we identified a pronounced expansion of ligand-gated ion channels (LGICs), whose primary role is to facilitate chemical transmission of signals to cells, and particularly neurons (Li et al. 2014). The expansion of LGICs, combined with the identification of three complex duplications in other gene families linked to neurofunction, suggest that *Homarus* lobsters have evolved additional chemosensory mechanisms for dealing with life in the benthos (Derby & Weissburg 2014). We also detected several expanded gene families linked to immunity, including: (i) pentaxins, an evolutionarily conserved family of proteins involved in acute immunological responses (Wang et al. 2020); (ii) immunoglobulin domains, which play a crucial role in the immune system; and, (iii) C-type lectins (identified as a simple duplication), which are known for their role in innate and adaptive antimicrobial immune responses (Brown et al. 2018). Although such families are often dynamic across many animal lineages, their significant expansion in *Homarus* nevertheless highlights candidate loci that represent promising targets for future investigation of lobster immunity, ecology, and evolution.

We also identified four complex duplications in gene families linked to genome organisation and several simple and complex duplications with putative functions in telomere maintenance, including: (i) reverse transcriptase (RNA-dependent DNA polymerase) which had six and seven copies in *H. gammarus* and *H. americanus*, respectively, and (ii) telomerase activating protein *est1*, identified as a simple duplication in *Homarus*, with both copies expressed in all *H. gammarus* tissues (Figure 3c). Previous research has found high telomerase activity across several lobster tissues (Klapper et al. 1998), suggesting that telomerase activation is a mechanism for maintaining long-term cell proliferation and senescence suppression in Homarid lobsters. The expansion of gene families linked to telomere maintenance and function may be a driver for high telomerase activity, which in turn could contribute to longevity. In addition, we detected several expanded gene families with functions in moulting, including chitin-related proteins, which may indicate a greater diversity of growth-related genes in *Homarus* relative to other decapods.

The identified simple duplications, such as those involving Lectin C-type domains and telomerase activating protein *est1*, are particularly informative because they represent clear candidates for lineage-specific duplication events that may have undergone functional divergence or neofunctionalisation during *Homarus* evolution. Future comparative analyses could assess whether these paralogs have acquired distinct functional specialisations, providing an opportunity to evaluate the adaptive significance of duplication in the *Homarus* lineage.

### Conclusions

We provide the first high-quality reference genome and annotation for the European lobster, together with a multi-tissue expression atlas (<u>LobsterGeneX</u>) to support genotype-to-phenotype studies. The assembly is highly complete but shorter than flow cytometry estimates, highlighting the repeat-rich nature of *Homarus* genomes and the need for future chromosome-scale scaffolding and repeat resolution. However, the discovery of a full-length *Dscam* locus demonstrates the value of this assembly for analysing complex, functionally important genes. Comparative analyses allowed us to date the divergence between *H. gammarus* and *H. americanus* to ∼26 Mya and identify *Homarus*-specific duplicate genes with functions in neurofunction, immunity, and telomere biology. Together, these resources lay a foundation for research into lobster immunity, longevity, adaptation and hybridisation, and will inform sustainable fisheries and aquaculture.

## Materials and Methods

### Genome sampling, DNA extraction and sequencing

Tissue for genome assembly was obtained from a wild adult male lobster (5–7 years old; carapace length >87 mm) collected in Cornwall, UK. Muscle tissue from the crusher claw was excised under sterile conditions and immediately flash-frozen in liquid nitrogen and stored at −80°C. High molecular weight (HMW) DNA was extracted using a modified salting-out protocol developed for lobster pleopods (Jenkins, Ellis & Stevens 2019). DNA was cleaned using the DNeasy PowerClean CleanUp kit (Qiagen, Germany) with modifications aimed at reducing DNA shearing: 30 μl of proteinase K was added to the tissue during the cell lysis step and incubated overnight in a thermomixer; for steps where vortexing was required, this approach was replaced by gently inverting samples 50 times, while precipitation of DNA using PowerClean columns was replaced with an isopropanol precipitation method.

Libraries were prepared using the Pacific Biosciences SMRTbell Express Template Preparation Kit (Pacific Biosciences, USA). Size selection was performed on a BluePippin (Sage Science, USA) at a >15 Kb cut-off threshold. Long-read sequencing was performed on a PacBio Sequel using diffusion loading across 14 SMRT Cells (1M v3 and Sequel Sequencing Chemistry v3) with a capture time of 10 h (106 Gb of data, approximating to ∼60X coverage). Using DNA obtained from the same sample, we performed Illumina short-read sequencing (600 Gb of data, approximating to 300x coverage) on two lanes of an Illumina NovaSeq S1 flow cell (150bp paired-end). Illumina data were cleaned and adaptors removed using fastp v0.23.4 (Chen et al. 2018).

### RNA sampling, extraction and sequencing

Four individual adult (two male and two female) individuals originating from wild populations collected from Weymouth (UK) were sampled for eye, gill, gut, heart, hepatopancreas, muscle, and nerve. Four individual adult females were sampled for ovary tissue, and four individual adult males were sampled for testes tissue. Four juvenile lobsters (moult stage 4, unknown sex) were obtained from the National Lobster Hatchery, Cornwall (UK). Individuals were placed on ice prior to tissue dissection. All tissues and organs were immediately snap-frozen in liquid nitrogen (juveniles as whole bodies) and stored at −80°C prior to sample preparation and RNA extraction.

All samples were disrupted by grinding frozen tissue fragments with liquid nitrogen before homogenisation with a rotor stator homogeniser in lysis reagent. RNA was extracted using the Qiagen mRNeasy Mini Kit (Qiagen, Germany), including an on-column DNase digestion step. RNA quality was measured using an Agilent 2100 Bioanalyzer with RNA 6000 nano kit (Agilent Technologies, CA, USA). ERCC Spike-in control mixes (Ambion via Life Technologies, Paisley, UK) were added to control for technical variation during sample preparation and sequencing and were analysed using manufacturer’s guidelines. mRNA isolation was performed via poly(A) enrichment using Tru-Seq Low Throughput protocol and reagents (Illumina, CA, USA). cDNA libraries were constructed using Illumina’s TruSeq Stranded mRNA Sample Preparation kit. Libraries were sequenced on an Illumina HiSeq 2500 (100 bp paired-end).

### Genome assembly

Genome size was estimated by *k*-mer profiling and flow cytometry. For *k*-mer profiling, the short-read Illumina data were *k*-merised into 19-, 21-, 23- and 25-mers using dsk v2.3.3 (Rizk et al. 2013)*. k*-mer profiles were assessed for genome size, heterozygosity and duplication rate in GenomeScope v2 (Ranallo-Benavidez et al. 2020). To assess the impact of repetitive sequences on genome size estimation, *k*-mers were summarised using coverage threshold cut-offs of 100,000 and 1,000,000. smudgeplot v0.2.3 was used to assess ploidy, counting heterozygous *k*-mer using kmc v3.0.0 (Kokot et al. 2017), using 21 and 1600 for the upper and lower cutoffs, respectively.

Genome size was estimated by flow cytometry using tissue derived from a single adult European lobster individual sampled from St Andrews (UK). Nuclei were isolated from eye tissue and two replicates of hepatopancreas. Mean nuclear fluorescence was measured on a Merck Milipore Muse Cell Analyzer flow cytometer (532 nm laser). Samples were calibrated against nuclei derived from European plaice (*Pleuronectes platessa*, 0.4 pg DNA diploid), European flat oyster (*Ostrea edulis*, 1.2 pg DNA diploid), giant tiger prawn (*Penaeus monodon*, 2.5 pg DNA diploid), and rainbow trout (*Oncorhynchus mykiss*, 2.5 pg DNA diploid, 3.5 pg DNA triploid). Genome size was calculated as the ratio of sample to standard mean fluorescence intensity multiplied by the known C-value of the standard.

We used Flye v2.8.1 (Kolmogorov et al. 2019) to assemble the PacBio data. Several parameters were initially explored, with the optimum assembly (based on N50 and the BUSCO completeness score) generated using the following parameters: (i) minimum read length of 1000 bp, (ii) --minimum-overlap 10000, and (iii) --asm-coverage 40. The assembly was polished using NextPolish v1.4.0 (Hu et al. 2019). For polishing, short-read data were aligned to the Flye assembly using bwa mem (Li 2013) and samtools v1.19 (Danecek et al. 2021) was used to correct (*fixmate*) and deduplicate (*markdup*) read pairings. Quality assessment after two rounds of polishing showed no improvement over the first round, so only one polishing iteration was used. Potential contamination in the assembly was assessed using BlobTools v1.1.1 (Laetsch & Blaxter 2017), identifying one potential contig as a contaminant, which was removed from the assembly. Haplotype purging was performed using purge_dups v1.2.5 (Guan et al. 2020). Genome completeness was assessed using BUSCO v5.7.1 (Manni et al. 2021) against the arthropoda_odb10 dataset (*n* = 1013) (Kriventseva et al. 2019). Merqury v1.3 (Rhie et al. 2020) was used to perform a *k*-mer assessment of quality and completeness using the short-read data (*k*-mer = 21).

### Genome annotation

Structural annotations were generated via the EMBL-EBI Ensembl Gene Annotation workflow (Aken et al. 2016). Following the Ensembl core database schema, the assembly was first screened for repetitive sequences using RepeatMasker v4.1.0 (Smit et al. 2015). A *de novo* repeat library was generated using RepeatModeller v2.0.1 (Flynn et al. 2020). However, because repeat modeller libraries may contain non-TE protein coding sequences, we opted to perform repeat library filtration of TEs using proteins from the sister species *Homarus americanus* (GCF_018991925.1) to avoid potentially over-masking protein coding loci with genuine TE-protein overlaps. After selecting the longest isoform per gene, the structural annotation was assessed using both BUSCO v5.7.1 (Manni et al. 2021) (arthropoda_odb10, *n* = 1,013) and OMArk v2.0.3 (Nevers et al. 2024) (Pancrustacea, *n* = 4,103).

### Gene expression atlas

For the tissue expression atlas “LobsterGeneX”, RNA-seq reads were aligned to the genome using STAR v2.7.3 (Dobin et al. 2012). Alignments were sorted and used as input to featureCounts (Liao et al. 2014), only counting gene features where both paired-end reads aligned. Normalised counts and the variance stabilising function (VST) were computed using DESeq2 (Love et al. 2014) and are available on the LobsterGeneX GitHub.

### Orthogroup identification

Searches for decapod proteomes were conducted with GoaT (Challis et al. 2023), using “Decapoda” as the query search. Initially, we downloaded and assessed the quality of 21 decapod proteomes. For each proteome, we selected the longest isoform for each protein-coding gene using AGAT v0.9.1 (Dainat 2024). The completeness of the filtered protein datasets was assessed with BUSCO (Manni et al. 2021) against the arthropoda_odb10 lineage database (Kriventseva et al. 2019) and the level of duplication was assessed with CD-hit v4.81 (Fu et al. 2012) using a clustering threshold of 0.8. To account for potential biases introduced by the different annotation pipelines used for *H. gammarus* and *H. americanus* (Prieto-Baños et al. 2025; Table 1), we also performed an annotation of *H. americanus* using the EMBL-EBI Ensembl Gene Annotation workflow. After quality assessment of the 21 initial protein datasets, we retained nine species for downstream analyses: *Homarus gammarus*, *Homarus americanus*, *Cherax destructor*, *Penaeus chinensis*, *Penaeus japonicus*, *Penaeus monodon*, *Penaeus vannamei*, *Portunus trituberculatus*, and *Procambarus clarkii*. We also included filtered proteomes for three outgroup species: *Daphnia magna*, *Drosophila melanogaster* and *Hyalella azteca*.

Orthology analysis followed previously published approaches (Regan et al. 2021). Quality-filtered proteomes were clustered into orthogroups using OrthoFinder v2.5.4 (Emms & Kelly 2019). OrthoFinder was run using Diamond v2.0.15 (Buchfink et al. 2021) for clustering, MUSCLE v3.8.1551 (Edgar 2004) for Multiple Sequence Alignment (MSA) and RAxML-NG v0.9.0 (Kozlov et al. 2019) for tree inference of the MSAs. Overall, 89.4% of genes were assigned to an orthogroup and 12.1% of orthogroups were species-specific.

### Divergence dating

To infer the divergence time between *H. gammarus* and *H. americanus*, we performed phylogenetic and divergence time estimation analyses. We restricted the taxon sampling to the nine Decapoda species used for orthogroup identification. Single-copy orthologs were aligned using MAFFT v7.505 (Katoh & Standley 2013). Poorly aligned regions were trimmed using trimAL v1.4.1 (Capella-Gutiérrez et al. 2009). A concatenated dataset of all 1295 single-copy orthologs was generated using AMAS (Borowiec 2016) and this was subsequently used to estimate a maximum-likelihood (ML) phylogeny using RAxML-ng v1.1.0 (Kozlov et al. 2019). We conducted 50 ML tree searches using 25 random and 25 parsimony-based starting trees using the LG+G4 amino acid substitution model. Following ML tree searches, we generated 500 nonparametric bootstrap replicates and checked for convergence post hoc using the --bsconverge command in RAxML-ng with a cutoff value of 0.03; we used the RAxML-ng --support command to compute bootstrap branch support values.

We used three calibration points: (1) at the root of the phylogeny (Crown Decapoda), (2) at Crown Astacidea, and (3) at the most recent common ancestor (MRCA) of *Penaeus vannamei*, *Penaeus monodon* and *Penaeus chinensis*. Calibration points were based on the posterior distributions of previous divergence time estimation analyses using fossil calibrations derived from more taxon-rich datasets (Bracken-Grissom & Ahyong 2014; Ma et al. 2009; Wolfe et al. 2019). To minimise the effects of among-lineage rate variation, we subsampled the 25 most clock-like single-copy orthologs using SortaDate (Smith et al., 2018). We conducted three divergence time analyses in Beast v2.6.7 (Bouckaert et al. 2019): (i) using a strict clock model and uniform priors for calibration points, (ii) using a strict clock model and normally distributed priors for calibration points, and (iii) using an uncorrelated log-normal relaxed clock model and normally distributed priors for calibration points. For each analysis, we executed three independent Markov chain Monte Carlo (MCMC) runs with different starting seed numbers. For each MCMC run, the best-fitted amino acid substitution model as inferred by ModelTest-NG (Darriba et al. 2019) was set for each partition, tree topology and clock model were linked by locus, and we set a Calibrated Yule model (Heled & Drummond 2012) as a tree prior. Each chain was run for 20 million generations, sampling every 20,000 generations and we checked each run for convergence of parameter values and age estimates by inspecting traces and effective sample sizes (ESS) in Tracer v1.7.1 (Rambaut et al. 2018). We then combined tree and log files from each set of three independent runs using LogCombiner v2.6.7 (Bouckaert et al. 2019) and generated maximum-clade-credibility trees with node heights set to mean age estimates discarding the first 10% of each MCMC as burn-in with TreeAnnotator v2.6.0 (Bouckaert et al. 2019). To assess the level of rate heterogeneity across lineages, we examined the coefficient of variation in clock rates. We also generated joint prior distributions for each of the three analyses by running MCMCs with no data.

### Identification of gene duplications and expansions in *Homarus*

To identify gene duplications and expansions specific to the *Homarus* branch, we compared the two *Homarus* species against seven decapod species and the three outgroup species listed above. We grouped *Homarus*-specific duplications into two categories: (i) simple duplications (a single duplication event in the *Homarus* branch), and (ii) multi-copy expansions (see below for definition). To explore the functional annotation of duplicated genes, we performed an analysis of the protein domains using InterProScan v.5 (Jones et al. 2014), using the top Pfam domain for functional inference.

We identified an initial set of 20 simple duplications and performed individual quality checks on these orthogroups to reduce the incorrect inference of split-gene annotations (*i.e.*, cases where a single gene is incorrectly annotated as two or more separate models) as gene duplicates. Protein sequences for each species for each orthogroup were aligned using MAFFT v.7.515 (Katoh & Standley 2013) and cleaned and trimmed using trimAl v.1.4.1 (Capella-Gutiérrez et al. 2009) for visual inspection in AliView v.1.27 (Larsson 2014). 12 candidate duplications were highly indicative of split-gene models and were removed, resulting in eight duplicated orthogroups for downstream analysis.

For multi-copy gene expansions, orthogroups were filtered for those showing a fold change > 2.0 and a Mann-Whitney-U p-value <=0.05 compared with the mean for all other Decapoda species. We only retained gene expansions which had an associated Pfam domain (96 orthogroups, comprising 203 total Pfam domains). Orthogroups present in less than five out of seven Decapoda species were subsequently removed, resulting in an analysis comprising 75 duplicated orthogroups.

## Data Availability

Raw sequence data are available under the BioProject accession PRJEB63315. The *Homarus gammarus* genome assembly is available on NCBI and the ENA under the accession GCA_958450375.1.

The Ensembl Metazoa annotation for *Homarus gammarus* is available at: https://metazoa.ensembl.org/Homarus_gammarus_gca958450375v1/.

The assembly is also available at the UCSC Genome Browser: https://genome.ucsc.edu/cgi-bin/hgTracks?genome=GCA_958450375.1&hubUrl=/gbdb/genark/GCA/958/450/375/GCA_958450375.1/hub.txt.

LobsterGeneX is freely available for use at www.LobsterGeneX.com. Expression matrices are available on the LobsterGeneX GitHub: https://github.com/Tom-Jenkins/LobsterGeneX/tree/main/public

