## Supplementary Figures for "A high-quality reference genome and tissue expression atlas for the European lobster (*Homarus gammarus*)"

**Figure S1:** smudgeplot analysis of ploidy levels

**Figure S2:** Characterisation of the *Dscam* gene

**Figure S3:** Gene expression of the *Dscam* gene

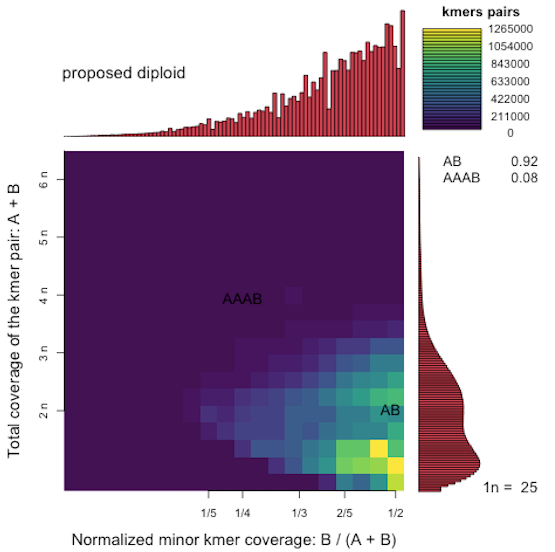

### **Figure S1.** smudgeplot analysis of ploidy levels.

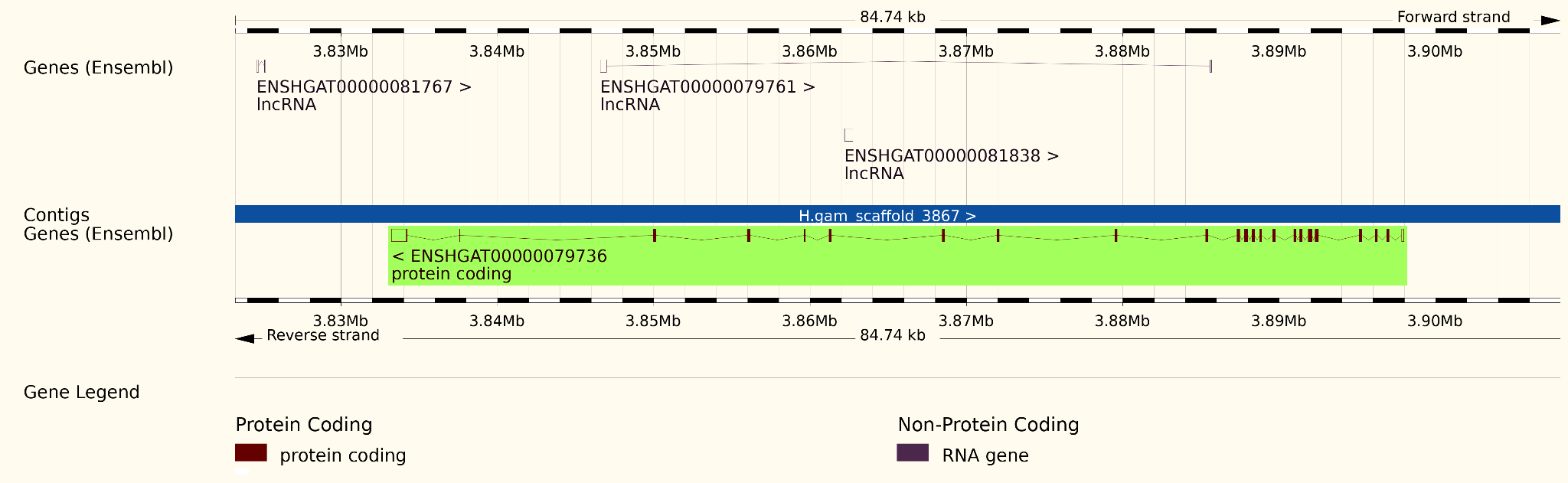

### **Figure S2. Characterisation of the *Dscam* gene (**[ENSHGAG00000010168](https://metazoa.ensembl.org/Homarus_gammarus_gca958450375v1/Gene/Summary?g=ENSHGAG00000010168;r=H.gam_scaffold_3867:3833226-3897963;t=ENSHGAT00000079736)) and of the transcript ENSHGAT00000079736 on scaffold_3867. This transcript has the Pfam domain: Down syndrome cell adhesion molecule (*Dscam*) C terminal.

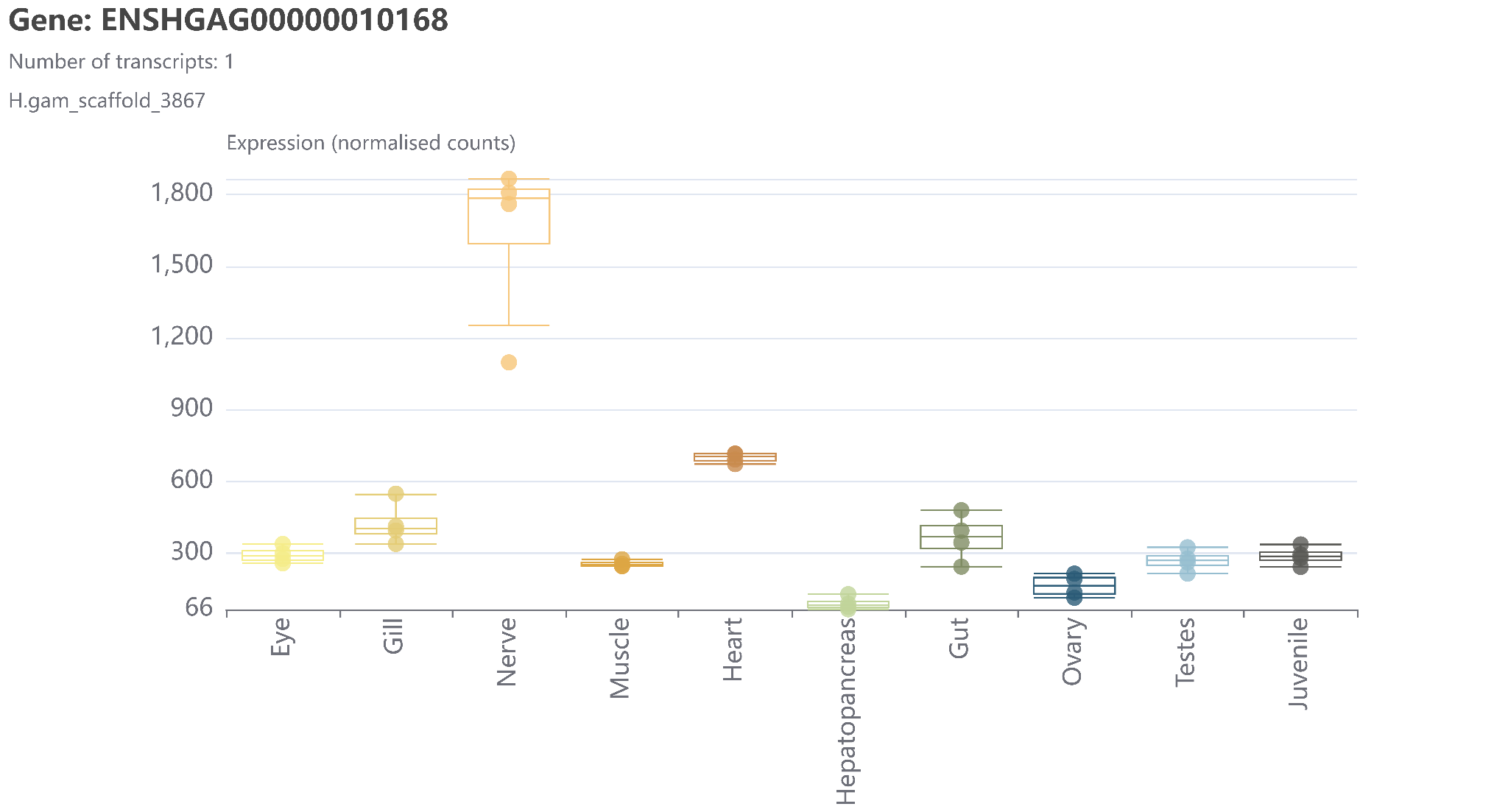

### **Figure S3.** Tissue expression profiles in the [ENSHGAG00000010168](http://enshgag00000010168) gene, generated via [LobsterGeneX](https://lobstergenex.com). This gene contains a transcript, ENSHGAT00000079736, that has the Pfam domain: Down syndrome cell adhesion molecule (*Dscam*) C terminal.
